# DModE: An end-to-end framework for Differential Modification and Expression Analysis of Nanopore direct RNA sequencing data

**DOI:** 10.64898/2026.06.09.731112

**Authors:** Johannes Miedema, Stefan Pastore, Luca Drescher, Alexander Haehnel, Anna Wierczeiko, Nicolò Alagna, Lioba Lehmann, Mark Helm, Tamer Butto, Susanne Gerber

**Author notes:** These authors contributed equally to this work (Johannes Miedema, Stefan Pastore).

## Abstract

**Summary:** Nanopore direct RNA sequencing (DRS) enables simultaneous quantification of transcript abundance and RNA modifications from native RNA molecules, providing a unique opportunity to study transcriptional and epitranscriptomic regulation within a single experiment. However, comprehensive analysis of DRS data remains challenging, as existing workflows typically focus on individual processing steps and often require manual integration of multiple software packages for expression analysis, modification detection, statistical testing, and visualization. Furthermore, integrated differential expression and differential RNA modification analysis at both gene and isoform resolution remains poorly supported by current workflows. Here, we present DModE (Differential Modification and Expression Analysis), an end-to-end framework for integrated analysis of Nanopore DRS data. DModE combines an Epi2ME-compatible Nextflow preprocessing workflow with a dedicated Python package for downstream statistical analysis, visualization, and reporting. The framework supports differential gene and isoform expression analysis, differential RNA modification analysis at genome and transcript level, metagene profiling, exploratory epitranscriptomic analyses, and integrated assessment of relationships between expression and modification dynamics. Results are automatically summarized in interactive HTML reports, facilitating reproducible and accessible data interpretation. By integrating transcriptomic and epitranscriptomic analyses within a single framework, DModE substantially simplifies comprehensive DRS data analysis and lowers the barrier for studying RNA modification biology using Nanopore sequencing.

**Availability and implementation:** The DModE Preprocessing pipeline is available on GitHub (https://github.com/johannesmiedema/DModE-preprocessing). The Python package can be installed via *pip* and is also available on GitHub (https://github.com/johannesmiedema/dmode).

## 1 Introduction

RNA modifications have emerged as a regulatory layer, influencing RNA stability, splicing, translation, and degradation. With over 150 chemically distinct modifications identified to date (Delaunay, Helm, and Frye 2024), their dynamic regulation is increasingly recognised as a fundamental layer of post-transcriptional control in both health and disease. Unlike short-read sequencing methods, DRS allows native RNA molecules to be sequenced directly, without reverse transcription or amplification, thereby preserving modifications as characteristic current deviations in the raw sequencing signal (Hewel *et al*. 2025). Notably, DRS simultaneously captures full-length transcript information, enabling quantitative analysis of both RNA modifications and gene expression - including isoform-level resolution - within a single experiment. While comprehensive workflows have been published for direct RNA modification analysis like MasterOfPores (Cozzuto *et al*. 2020) or nanoseq (https://github.com/nf-core/nanoseq), these tools mostly focus on separated analysis of gene expression or RNA modification and often require substantial manual integration of multiple software packages for comprehensive downstream analysis. Furthermore, most RNA modification pipelines do not consider isoform-specific patterns. Thus, the integrated analysis of transcriptomic and epitranscriptomic features in full-length isoforms is cumbersome and lacks a unified framework. To our knowledge, no currently available workflow provides integrated differential gene expression and differential RNA modification analysis at both genome and isoform resolution within a single framework.

To address this gap, we present DModE (Differential Modification and Expression Analysis), a comprehensive framework that combines preprocessing and downstream statistical analysis of DRS data into a single, unified framework. DModE is distributed as an Epi2ME-compatible Nextflow pipeline consisting of a preprocessing module and a centralized Python package for statistical analysis and visualization, enabling reproducible, user-friendly analysis and interactive HTML reports directly from raw sequencing data (pod5 files), making comprehensive DRS analysis accessible to a broad spectrum of researchers.

## 2 Methods

### 2.1 Usage

DModE is structured as a two-stage framework (Fig. 1). The first stage processes the raw input files into count tables. A Nextflow-based preprocessing pipeline performs basecalling and modification calling from raw pod5 files. Basecalled sequences are aligned to the genome and transcriptome, and the resulting BAM files are used for quantification of read abundance and modification on gene and transcript level. Outputs are stored as count tables and per-sample modification pileup BED files. The second stage performs various downstream analysis and data visualization steps. The DModE Python package integrates these outputs and performs quality control (QC), differential expression analysis and differential modification analysis. The results are exported in an HTML report of high-quality figures and summary tables. The two components are designed to be used sequentially but can also be run independently, allowing users who have already completed preprocessing with separate tools to enter the workflow at the statistical analysis stage.

**Figure 1:**
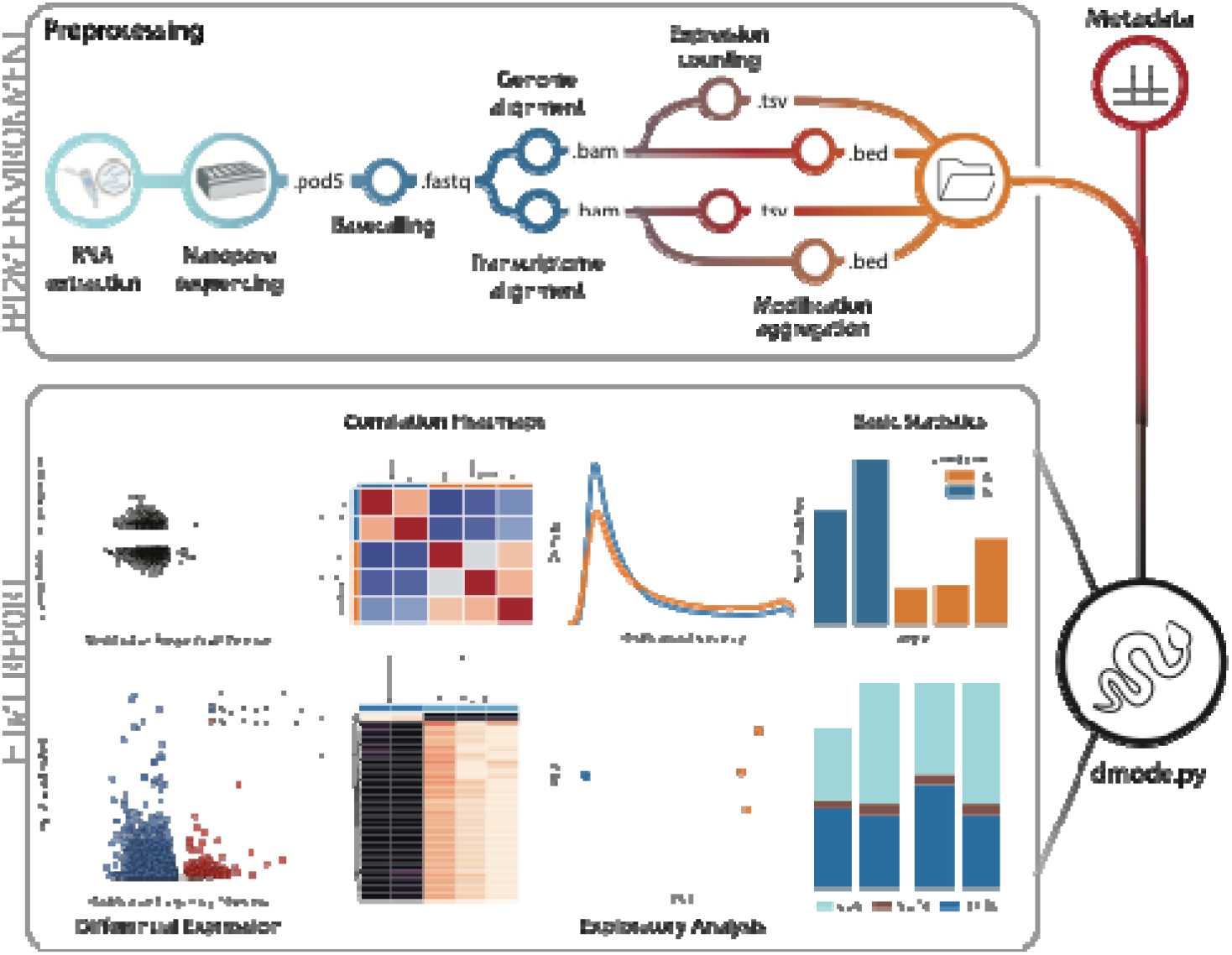
Schematic overview of the DModE framework. The first part consists of a Nextflow preprocessing pipeline, which can be run within the Epi2ME environment with a graphical user interface. The second part of the framework utilizes the DModE Python package (API and CLI) for statistical downstream analyses, with the results presented in interactive and user-friendly HTML reports.

### 2.2 Preprocessing pipeline

The DModE preprocessing pipeline is implemented in Nextflow (Di Tommaso *et al*. 2017) and is fully compatible with the Epi2Me platform, providing a graphical interface for users who prefer not to work on the command line. All processes are containerised using Docker (Merkel 2014), ensuring reproducibility across computing environments. The pipeline is launched from a single command pointing to a directory of POD5 files and the required reference files (genome FASTA, transcriptome FASTA, and GTF annotation).

The pipeline executes the following steps for each sample. First, raw POD5 signal data are basecalled using Dorado (https://github.com/nanoporetech/dorado), supported modifications include N6-methyladenosine (m6A), 5-methylcytosine (m5C), pseudouridine (pseU), inosine, and 2’-O-methylated nucleosides (Am, Cm, Um, Gm). Second, basecalled reads are aligned to both the reference genome and the reference transcriptome using Minimap2 (Li 2018). Third, gene-level expression counts are generated from genome-aligned BAM files using FeatureCounts (Liao, Smyth, and Shi 2014), while isoform-level transcript quantification is performed on transcriptome-aligned BAM files using Salmon (Patro *et al*. 2017). Fourth, QC statistics are generated per sample using NanoComp (De Coster and Rademakers 2023), comparing key QC metrics between genome and transcriptome alignments. Finally, Modkit (https://github.com/nanoporetech/modkit) pileup is run on both the genome-aligned and transcriptome-aligned BAM files to produce per-position modification frequency tables in BED format. Modkit confidence thresholds are fully configurable by the user. The pipeline produces structured output directories for each analysis step, which can be directly used by the DModE Python package.

### 2.3 Statistical analysis and visualisation

The DModE Python package provides a unified interface for downstream analysis, accepting the structured outputs of the preprocessing pipeline. Prior to differential analysis, QC and exploratory visualisations can be generated on the epitranscriptomic level. Those include Principal component analysis (PCA, Supplementary Fig 1B), Uniform Manifold Approximation and Projection (UMAP) plots from modification frequency tables, sample-to-sample correlation heatmaps (Supplementary Fig 1A), per-sample modification site barplots (Supplementary Fig 1C), frequency distribution and violin plots (Supplementary Fig 1D), metagene modification profiles (Supplementary Fig 2), and intersection barplots for cross-referencing with known modification databases. The DModE package can be used both as command line interface (CLI) or as application programming interface (API).

Differential gene expression and transcript expression analyses are performed using DESeq2 via the pydeseq2 Python package (Muzellec *et al*. 2022), accepting count tables from either FeatureCounts (genome level) or Salmon (isoform level). Differential RNA modification analysis can be performed at both the genome and transcript level. At each position, statistical significance is assessed using Fisher’s exact test for single-sample comparisons or logistic regression when multiple samples per condition are available. A sliding linear model (SLIM) method or Benjamini-Hochberg correction is applied to correct for the False Discovery Rate (FDR) in multiple hypothesis testing. The statistical framework was reimplemented and adapted in Python (Supplementary Fig 3) from the R package methylKit (Akalin *et al*. 2012).

Coverage and modification frequency filters can be applied prior to testing, and significant positions are annotated via GTF lookup and summarised at gene or transcript level. Furthermore, DModE also connects differential expression and modification data within a single visualisation, enabling users to rapidly discern whether genes with significant modification changes are also transcriptionally dysregulated (Supplementary Fig. 4).

### 2.4 Installation and requirements

The DModE preprocessing pipeline requires Nextflow and Docker. All containers are pre-configured in the pipeline and pulled automatically at runtime. Recommended hardware for preprocessing is 32 CPU cores and 64 GB RAM; the minimum specification is 16 cores and 32 GB RAM. For Basecalling, a NVIDIA GPU is required. Full installation instructions and documentation are available at https://github.com/johannesmiedema/DModE-preprocessing.

The DModE Python package requires Python 3.10 and is installable via pip (*pip install dmode*). Full installation instructions and documentation are available at https://github.com/johannesmiedema/dmode. Moreover, the github repository includes an application tutorial (https://johannesmiedema.github.io/dmode/), which is based on a publicly available test dataset comprising 2 replicates of UHRR and 3 replicates of HEK293T cells (Hewel *et al*. 2025) The test dataset has been preprocessed using the Nextflow based DModE pipeline to simplify the testing of the CLI/API based DModE downstream analysis. The preprocessed test data is available at Zenodo (https://zenodo.org/records/20496900).

## 3 Discussion

DModE provides a comprehensive, end-to-end framework for the analysis of DRS data, addressing a major limitation of current workflows, which require manual integration of multiple software packages across preprocessing, statistical analysis, expression analysis, and RNA modification analysis. By combining these analytical layers within a unified, streamlined pipeline, DModE substantially lowers the technical barrier to investigate transcriptomic and epitranscriptomic changes simultaneously. The Epi2ME integration further broadens accessibility by providing a graphical user interface for the preprocessing module.

A key strength of DModE is the integrated analysis of RNA modifications and gene expression at both genome and isoform resolution. This feature enables researchers to directly investigate relationships between transcriptional and epitranscriptomic regulation within a single analytical workflow, facilitating biological interpretation of DRS datasets. As DRS continues to mature, DModE provides a flexible foundation for integrated transcriptomic and epitranscriptomic data analysis and opens new possibilities for studying the functional consequences of epitranscriptomic changes.

## Author contributions

S.G. supervised the project. J.M. and S.P conceived the study, J.M., S.P. and A.W. designed, implemented and tested the DModE python package. J.M., S.P. and L.D. designed, implemented and tested the preprocessing pipeline. S.P. and A.H. implemented the API of the DModE Python package. J.M. and S.P. wrote the manuscript. S.G. N.A., L.L., M.H. and T.B. edited the manuscript and provided valuable input and feedback in various discussions.

## Conflict of interest

None declared

## Funding

This work was funded by Deutsche Forschungsgemeinschaft (DFG, German Research Foundation, project no. 439669440 TRR319 RMaP TP A07 and C04 (S.G.) and C01 (M.H). S.G. and L.L. acknowledge funding from the Boehringer Ingelheim Stiftung.

## Notes

### Competing Interest Statement

The authors have declared no competing interest.

