## Supplemental Information for "DModE: An end-to-end framework for Differential Modification and Expression Analysis of Nanopore direct RNA sequencing data"

^#^Corresponding Author

**Test dataset**

A test dataset of (Hewel *et al.* 2025) comprising 2 replicates of the Universal Human Reference RNA (UHRR) and 3 replicates of RNA derived from Human Embryonic Kidney (HEK293T) cells transfected with artificial snoRNAs was used to demonstrate the functionality of DModE. UHRR RNA is extracted from several tumor cell lines (Novoradovskaya *et al.* 2004). RNA of HEK293T cells is derived from human embryonic kidney tissue, which is not of malignant origin. However, transformation, immortalization and extensive passaging of HEK293T cells have let to the expression of proteins which are not found in the bona fide celltype (Kuadkitkan, Wikan, and Smith 2016). Nevertheless, a strong deviation on the modification and gene expression landscape of cancer related genes in UHRR is expected compared to HEK293T. Raw pod5 data was obtained from the European Nucleotide Archive at EMBL-EBI under the accession number PRJEB74238. Raw data was preprocessed using the DModE-preprocessing pipeline (<https://github.com/johannesmiedema/DModE-preprocessing>). Subsequent statistical analyses, including generation of the interactive HTML report was performed using the DModE Python package (<https://github.com/johannesmiedema/dmode>). All plots can be reproduced following the DModE tutorial ([https://johannesmiedema.github.io/dmode/)](https://johannesmiedema.github.io/dmode/)

**Nextflow based preprocessing pipeline**

Preprocessing pipeline reference data has been retrieved from GENCODE (Mudge *et al.* 2025). The primary assembly fasta for the genome, the transcript sequences for the transcriptome and the comprehensive GTF file of hg38 v49 have been downloaded. Preprocessing was started with ONT based raw current files in pod5 format. Reads were basecalled with dorado v1.3.0 using the super accuracy model v5.2.0 with m5C_2OmeC, inosine_m6A_2OmeA, pseU_2OmeU, 2OmeG v1 modification calling models. Resulting reads in unaligned BAM format were aligned with minimap2 (v2.28) to genome in splice-ont mode and to transcriptome in map-ont mode including MD flags. Nanocomp (v0.6.0) was performed for quality control. For the assessment of modification abundancies, modkit (v0.6.1) was used on genome and transcriptome level. To the dataset of Hewel and Wierczeiko et al. 2025 we used a mod-threshold of 0.98 on each modification and a filter-threshold of 0.8 for canoncial bases.

Transcript quantification was based on transcriptome aligned bam files using salmon (v1.10.3) with minAssignedFrags 0. Quantification of aligning to the genome was performed using the FeatureCount module of the subread package (v2.0.2) in long read mode (-L).

**Downstream analysis using DModE**

In the following section we highlight some analytical opportunities, using DModE. Plots generated in the following sections were created on m6A sites only and genome level only to showcase its functionality.

**Basic statistics**

Pairwise correlation of datasets by modification frequencies or Principal Component Analysis (PCA) of modification frequency revealed similarity in modification composition of datasets. Samples of the same condition show high similarity and are clearly separable from samples of the other condition. Plots of modification frequency distributions and the absolute number of modified sites delivered comprehensive insight into the dataset composition and the difference in frequency dynamics between HEK293T and UHRR cells. (Sup Fig.1).

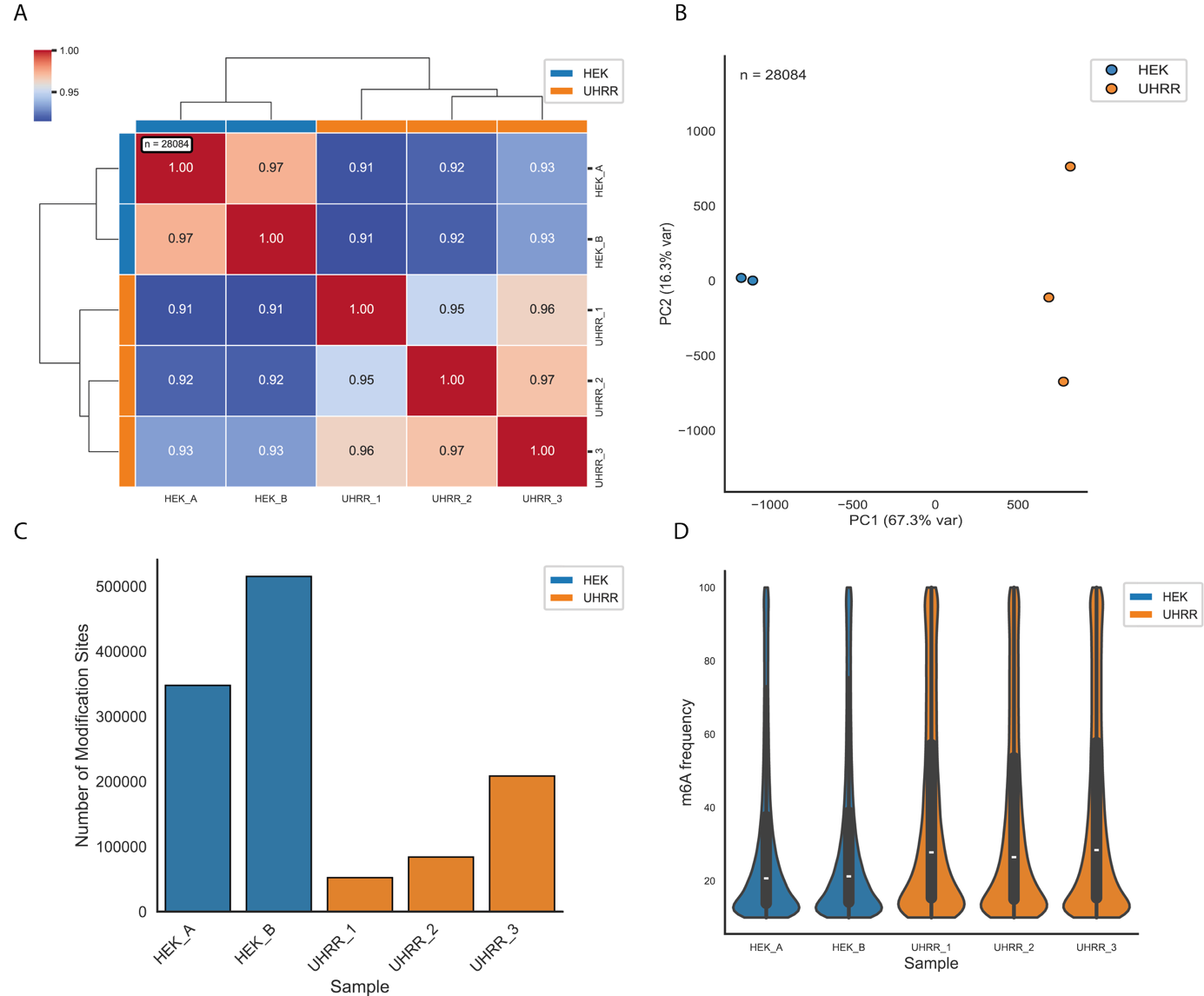

Supplementary Figure 1: **A)** Pairwise Pearson correlation of m6A modification frequencies of different datasets. **B**) Principal component analysis based on modification frequencies. **C)** Absolute modification abundance per dataset. **D)** Violinplots of modification frequency distribution per dataset.

**Metagene plots**

*Gene Body Model Construction*

To map modification sites onto a standardized transcript coordinate system, we first constructed a reference gene body model from a GTF annotation. Gene, transcript, CDS, and UTR features were extracted, and feature lengths computed from genomic coordinates. For each protein-coding gene, we selected a single representative transcript using a hierarchical scheme: the MANE Select transcript where available, falling back to the Ensembl canonical transcript, and otherwise to the first annotated transcript of the gene.

For each selected transcript, CDS and UTR segments were retained and the untranslated regions were assigned 5′ or 3′ identity based on their genomic position relative to the coding sequence, with the assignment inverted for genes on the minus strand to preserve biological 5′→3′ orientation. Features were ordered along the transcript and assigned cumulative transcript-relative start and end coordinates, yielding a contiguous gene body model spanning the 5′UTR, CDS, and 3′UTR.

*Modification Site Assignment*

Per-sample modification calls (modkit bedMethyl output) were intersected with the gene body model using genomic interval overlap joins, associating each modification site with its host transcript feature and recording validity, no-call, and failure read counts as coverage measures. Sites were optionally filtered by a minimum coverage threshold and could be restricted to a specific gene or modification type. Modification positions and feature boundaries were converted to transcript-relative coordinates, with positions on minus-strand genes reflected so that all transcripts share a common 5′→3′ frame.

*Metagene Profile Generation*

Each transcript region was scaled to a fixed number of bins (5′UTR, CDS, and 3′UTR scaled independently), producing a normalized metagene coordinate of fixed total length regardless of native transcript length. Each modification site was binned by its relative position within its assigned region, and the underlying read coverage was tallied in parallel.

Two binning modes were implemented. In the inclusive mode, positions were mapped directly onto the scaled coordinate, retaining intronic spacing. In the exclusive mode, only exonic intervals were extracted, concatenated, and linearly interpolated back to the fixed bin number, removing intronic contributions and yielding a purely exonic profile.

Per-transcript count and coverage vectors were normalized (by maximum, sum, or mean) and averaged across all transcripts within each experimental condition to produce condition-level metagene profiles. These profiles were plotted along the concatenated 5′UTR–CDS–3′UTR axis, with region boundaries demarcated, showing both relative modification abundance and coverage.

It should be emphasized that coverage is recorded only at positions that carry a called modification, since the intersection retains exclusively those sites present in the modification call set. The resulting coverage track therefore does not represent transcriptome-wide read depth, but rather the depth observed specifically at modified positions. Where a single genomic position gives rise to multiple modification calls of different types, coverage is counted only once per position to avoid inflating the signal through duplicate entries. Consequently, the coverage profiles should be interpreted as the read support underlying detected modification sites, and not as a general measure of expression or sequencing depth along the transcript. Regions devoid of called modifications contribute no coverage information, and apparent differences in the coverage track may reflect the distribution of modification sites themselves rather than true variation in underlying read depth.

*Feature-Level Summaries*

In addition to positional profiles, the distribution of modification sites across transcript features was summarized. Unique modification sites were counted per feature for each sample and aggregated by condition, then visualized as absolute and relative stacked bar plots at both the sample and condition level.

Metagene plots for m6A sites of UHRR and HEK293T cells (Sup. Fig. 2) revealed similar distribution patterns as shown in previous publications (Tegowski, Flamand, and Meyer 2022, Liu *et al.* 2023).

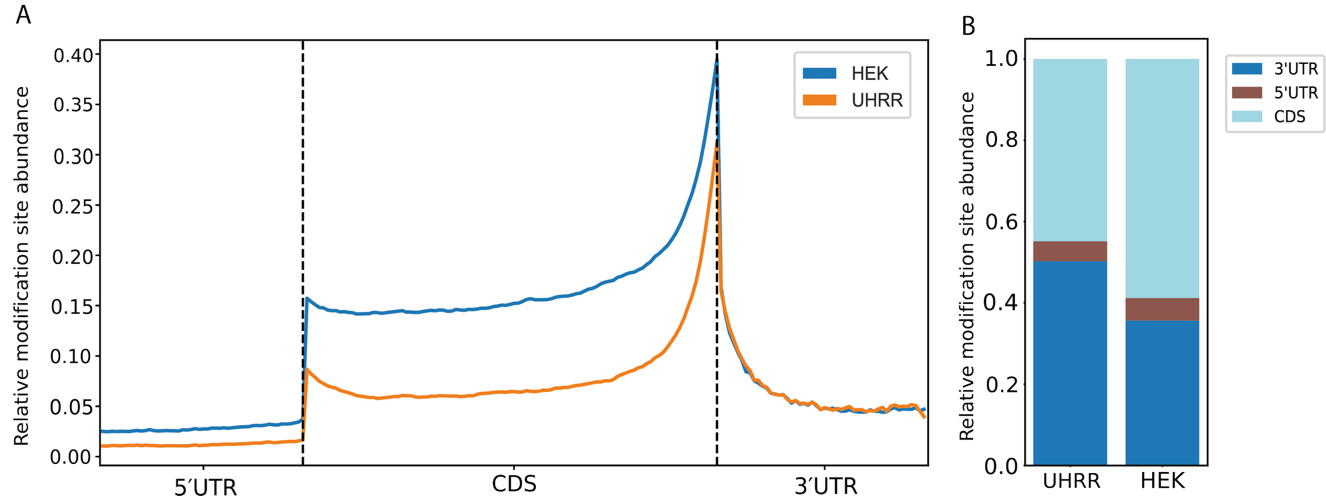

Supplementary Figure 2: **A)** m6A based metagene plot on UHRR and HEK293T samples. **B)** Relative modification site abundance on 5'UTR, CDS and 3'UTR.

**Differential expression and modification analysis**

*Comparison of Differential modification module with MethylKit*

Differential modification analysis was implemented according to the logics of Methylkit (v0.99-2). We therefore performed a comparative analysis on the m6A sites of the UHRR vs. HEK293T dataset using Methylkit and DModE. The results of both tools revealed great alignment with 14668 shared significant differential modified sites out of 38297 modified sites in total and an adjusted p-value based Pearson and Spearman correlation coefficient of one. A set of 25 significant modified sites was only detected in MethylKit, which could be explained by minor numeric inconsistencies of the computed p-adjusted value around the 0.05 significance boundary, likely caused by technical differences in the implementation of the different programming languages. (Sup. Fig.3)

While differential modification analysis was based on the methodology of MethylKit, differential expression analysis was performed using PyDESeq2 (v0.5.4). As with the correlation analysis of modification frequencies (Sup. Fig. 1), sample-to-sample comparisons of differential expression demonstrated high consistency within conditions (Sup. Fig. 4A). The top 50 differentially expressed genes included a variety of oncogenes overrepresented in UHRR. Similarly, strongly significant differentially modified sites (p <= 0.0001) were found to reside on oncogenes. (Sup. Fig. 4A, Sup. Fig. 4B, Sup. Fig. 4D) To investigate potential interdependencies between gene expression and modification, significant differential expressed genes and significant differential modified sites were investigated on their direction of difference and potential correlative properties. Assuming a linear Pearson correlation, the analysis did not reveal correlations in either direction across the modification and gene expression landscape. Rank based Spearman correlation could be shown for upregulated modification frequencies compared to differential expressed genes with coefficients of p=0.8 (expr. Up, mod. freq. up) and p=0.55 (expr. down, mod. freq. up). GO term analysis using the DisGeNET database (Piñero *et al.* 2017) applied to genes exhibiting significant differences in both modification frequency and gene expression, predominantly returned cancer-related terms, consistent with the cancer-derived origin of the UHRR Cell line (Sup. Fig. 4E).

Supplementary Figure 3: **A)** Correlation of adjusted pvalues assessing significance for differential modification analysis between MethylKit and DModE on m6A sites of the UHRR vs HEK293T dataset. **B)** Adjusted pvalues close to the 0.05 significance boundary, revealing a minor numeric inconsistency between results of MethylKit and DModE, which can be the result of technical differences of the programming languages.

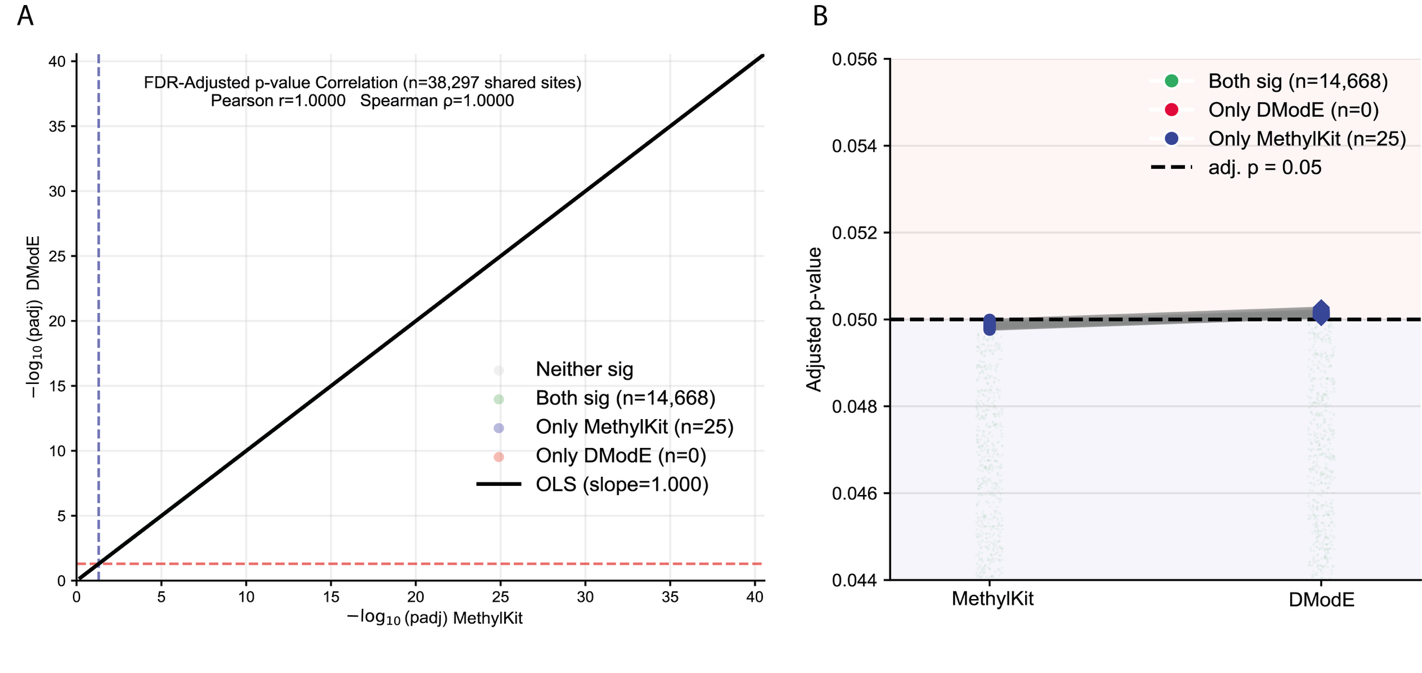

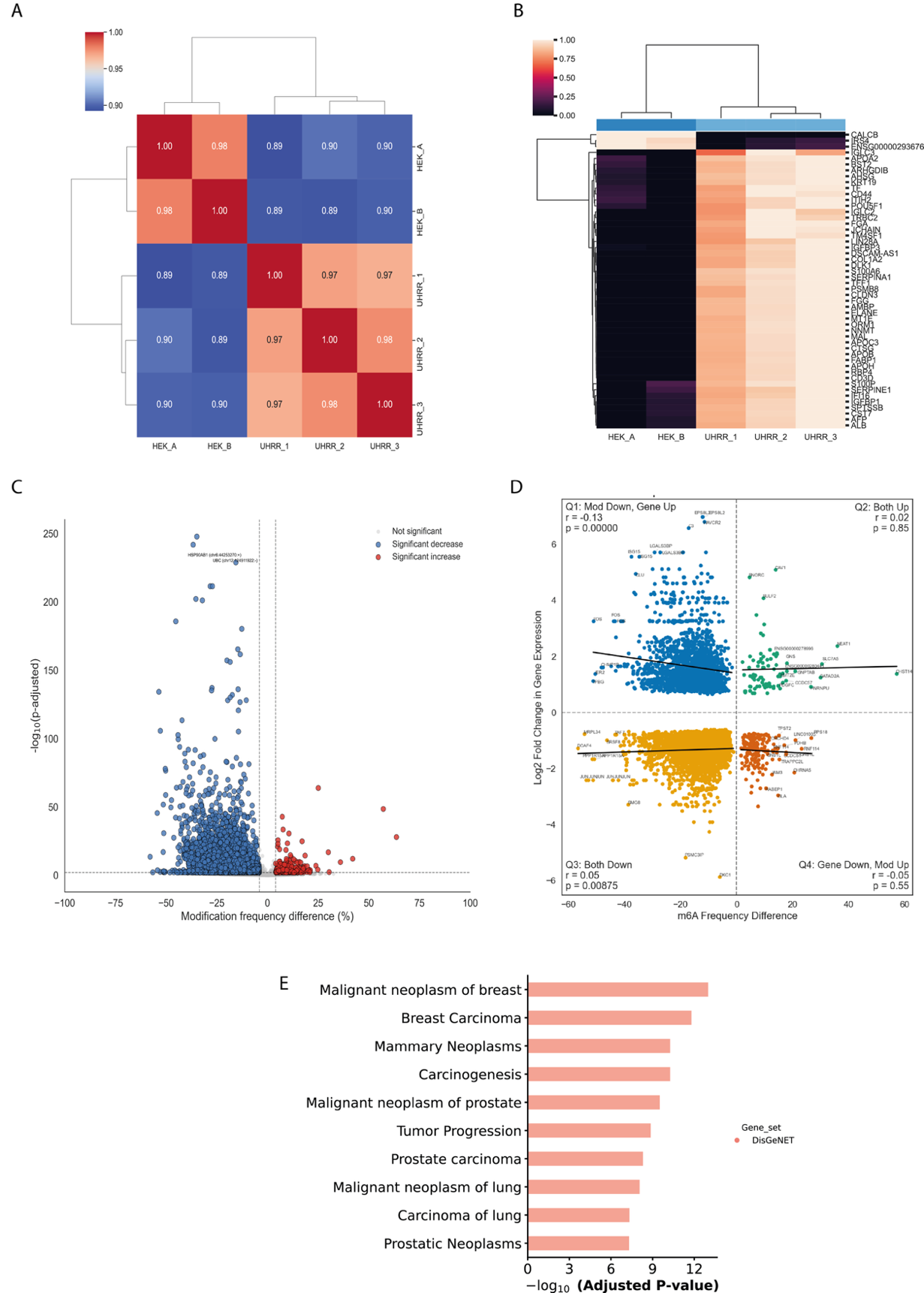

Supplementary Figure 4: **A)** Sample to sample plot with Pearson correlated logarithmic gene counts between UHRR and HEK293T samples. **B)** Heatmap of top50 differential expressed genes in UHRR vs HEK293T cells. **C)** Volcano plot of differential modified m^6^A sites showing modification frequency differences compared to the negative logarithm of p-adjusted- **D)** Quadrant plot comparing modification frequency difference with Log2FoldChange expression. Pearson and Spearman correlation coefficients are computed. Only significant modified m^6^A sites and significant differential expressed genes are shown.
